# Vaccine Elicitation of HIV Broadly Neutralizing Antibodies from Engineered B cells

**DOI:** 10.1101/2020.03.17.989699

**Authors:** Deli Huang, Jenny Tuyet Tran, Alex Olson, Thomas Vollbrecht, Mariia V. Guryleva, Mary Tenuta, Roberta P. Fuller, Torben Schiffner, Justin R. Abadejos, Lauren Couvrette, Tanya R. Blane, Karen Saye, Wenjuan Li, Elise Landais, Alicia Gonzalez-Martin, William Schief, Ben Murrell, Dennis R. Burton, David Nemazee, James E. Voss

## Abstract

HIV broadly neutralizing antibodies (bnAbs) can suppress viremia and protect against infection^1^. However, their elicitation is made difficult by low frequencies of appropriate precursor B cell receptors and the complex maturation pathways required to generate bnAbs from these precursors^2^. Antibody genes can be engineered into B cells for expression as both a functional receptor on cell surfaces and as secreted antibody^3–5^. Here, we show that HIV bnAb-engineered primary mouse B cells can be adoptively transferred and vaccinated in immunocompetent wild-type animals resulting in the expansion of durable bnAb memory and long-lived plasma cells. Somatic hypermutation after immunization indicated that engineered cells have the capacity to respond to an evolving pathogen. These results encourage further exploration of engineered B cell vaccines as a strategy for durable elicitation of HIV bnAbs to protect against infection and as a contributor to a functional cure.

---

Critical support for efforts to design an HIV vaccine that would elicit broadly neutralizing antibodies (bnAbs) comes from experiments that show such antibodies are capable of providing sterilizing immunity against clinically relevant (so-called Tier 2) virus challenge in macaque and humanized mouse models^6–8^. Such antibodies are difficult to elicit through vaccination due in part to genetic limitations of the human antibody repertoire. Characterization of a variety of bnAbs discovered in chronically infected patients shows that they generally derive from B cell precursors with uncommon antigen receptor features such as long third heavy chain complementarity determining regions (CDRH3s) which then require extensive somatic hypermutation^8,9^.

The gene sequences of HIV bnAbs and other desirable antibodies can however be engineered into the genomes of *ex vivo* activated primary B cells, such that they are expressed as functional B cell antigen receptors (BCRs) using endogenous heavy chain (HC) constant genes^3–5^. As such, engineered BCRs can undergo class switching for eventual secretion as protective antibodies from plasma cells. B cells engineered in this way have been shown to confer protective levels of pathogen-specific antibody *in vivo* for several weeks following adoptive transfer into immunocompromised hosts^5^, or for several days following transfer into immunocompetent hosts^4,5^.

Long-term expression of HIV bnAbs generated from engineered B cells, which could be boosted through vaccination and which could mature in affinity against relevant viral sequences in immunocompetent hosts, has potential as an attractive functional HIV cure strategy, given that bnAbs administered passively in the context of infection have been shown to suppress viremia, kill infected cells, and enhance host immunity^1,10,11^. Wild-type (WT) black 6 (C57BL/6J) mice represent a useful model in which such an engineered B cell vaccine could be developed because of similarities between mouse and human humoral immune systems, and because the mouse antibody repertoire is also strongly genetically restricted in its ability to elicit HIV bnAbs^12–14^. In activated primary B cells, we used CRISPR-Cas9 to insert the VRC01 HIV bnAb^15,16^ light chain (LC) and HC variable region (VDJ) into the mouse HC locus at *J4* using a homology directed repair (HDR) genome editing strategy. The inserted VRC01 gene is expressed under the control of a HC V-gene promoter as a single mRNA, which is post-transcriptionally spliced to endogenous HC constant genes. A P2A self-cleaving peptide sequence downstream of the mouse kappa (κ) constant gene separates the VRC01 light and heavy chains, allowing them to pair and form a functional cell surface expressed BCR (*H-targeting*) (Fig. 1**a**). We chose to use the VRC01 genes because this prototype CD4 binding-site bnAb, which blocks entry of the virus into target CD4^+^ T cells, has been extensively tested in the clinic for its ability to suppress viremia in patients and prevent infection after administration as a recombinant monoclonal antibody^17–19^ (clinicaltrials.gov NCT02568215, NCT02716675).

**Figure 1:**
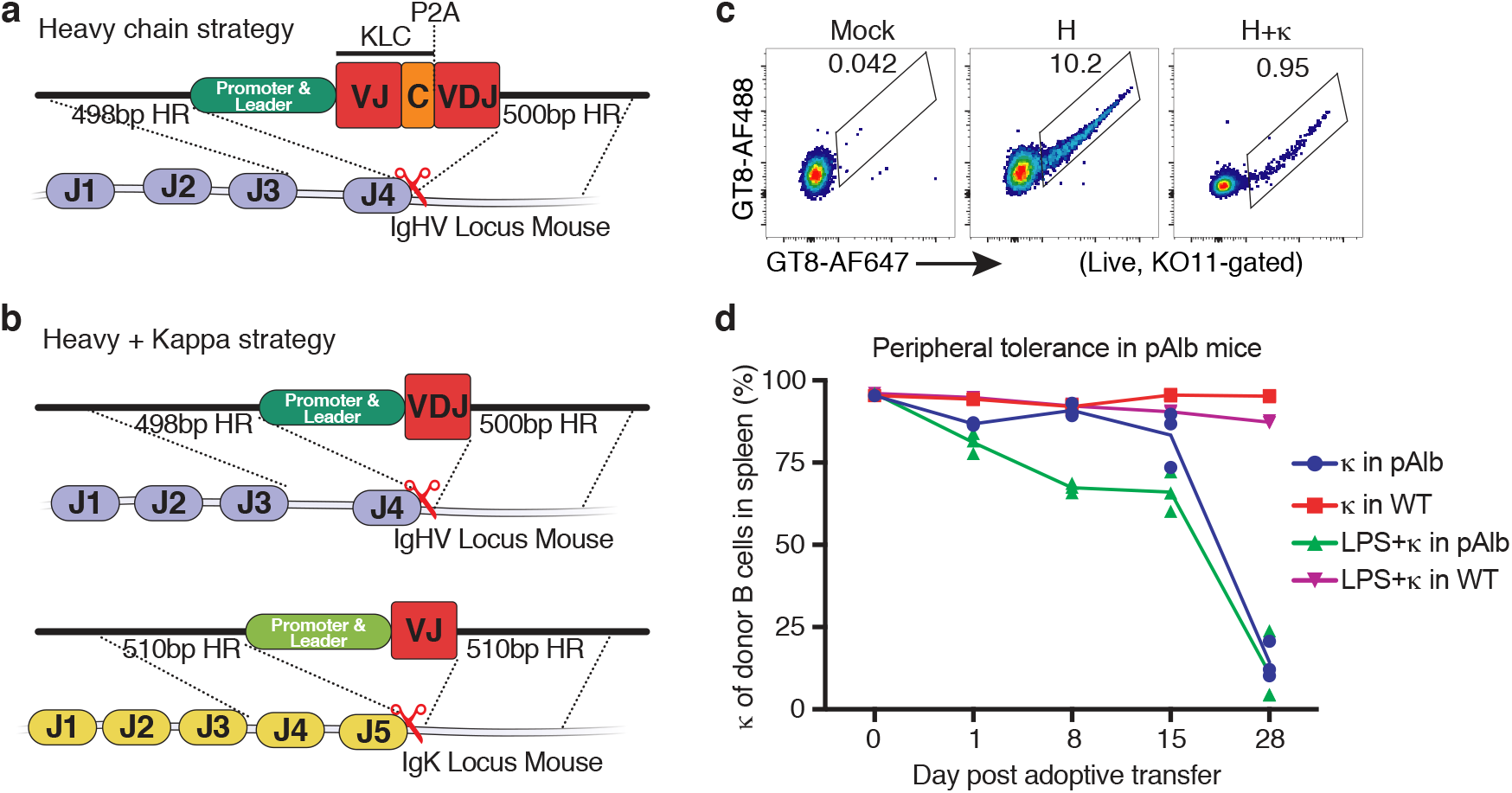
Primary B cell engineering and engraftment. **a, Targeting antibody genes to the mouse heavy chain (HC) locus (*H-targeting*).** Donor DNA encoding: a HC V-gene promoter, the VRC01 kappa chain (with mouse constant gene and P2A peptide), the VRC01 HC variable (VDJ) region, and constant gene donor splice site is inserted at a CRISPR-Cas9 cut site in the *IgH-J4* gene for expression as a functional antigen receptor using endogenous downstream HC constant genes. Regions of homology (HR) flanking bnAb donor DNA allow for its incorporation at the DNA break-site by homology directed repair (HDR). **b, Targeting antibody genes to the mouse heavy and kappa loci (*H+κ targeting*).** To engineer the *IgH* locus, donor DNA encoding: a HC V-gene promoter, VRC01 VDJ gene, and constant gene donor splice site is inserted as above. To engineer the Igκ locus, donor DNA encoding: a κ V-gene promoter, VRC01 κ variable (VJ) region, and constant gene donor splice site is likewise inserted via a CRISPR-Cas9 cut site in the *IgK-J5* for expression of VRC01 H and κ chains from their endogenous loci spliced to cell-native constant genes. **c, Targeting efficiency.** The VRC01 specific eOD-GT8 immunogen was used as a probe to identify successfully targeted cells by flow cytometry. Conditions depicted include only LPS activated (Mock), *H+κ*, and *H-targeted* B cell cultures. **d, Auto-reactive B cells are deleted *in vivo* after LPS culture and adoptive transfer.** Untouched and LPS activated B cells were transferred to pAlb mice expressing anti-kappa super-antigen in the liver. The fraction of donor B cells which were κ^+^ was analyzed at the indicated times post-cell transfer. Data points represent the average values from n=3 animals.

A second strategy was also employed, in which the VRC01 heavy and kappa variable genes were targeted separately to their endogenous loci for expression from V-gene promoter-controlled transcripts spliced to cell-native H or κ constant genes (*H+κ targeting*) (Fig. 1**b**). Targeting success was monitored by detecting cell surface expressed VRC01 using flow cytometry probes specific for this antibody, eOD-GT8 and KO11^20,21^. Targeting by *H* or *H+κ* strategies routinely resulted in engineering efficiencies of 10% and 1%, respectively (Fig. 1**c**).

Current engineering strategies can result in the expression of self-reactive BCRs due to pairing of engineered Ig chains with those endogenously produced by the targeted cell^3–5^. While tolerance mechanisms in the periphery of mice and humans generally ensure that autoreactive B cells are non-functional^22–25^, we sought to ensure that this would still be the case for cells activated *ex vivo* using the methods we required for efficient HDR based B cell genome editing. To do this, we made use of transgenic mice expressing an Igκ chain-reactive super-antigen on the surface of hepatocytes (pAlb mice)^26^. B cells purified from the spleens of WT donor animals by negative selection were adoptively transferred into host mice directly, or after *ex vivo* activation using the toll-like receptor 4 (TLR4) agonist, lipopolysaccharide (LPS). When transferred as non-self-reactive B cells into WT mice, Igκ^+^ cells survived. When transferred into pAlb mice, the now self-reactive Igκ^+^ cells were deleted as expected for both directly transferred and *ex vivo* activated cells (Fig. 1**d**). These results suggest that auto-reactive B cells generated during the engineering step will remain subject to peripheral tolerance mechanisms *in vivo*.

B cells purified from the spleens of WT donor animals by negative selection were then activated with LPS, targeted, and transferred into WT recipients to assess their status *in vivo*. After 14 days, donor cells that were directly transferred had low frequencies of memory B cells (MBCs), whereas 70-80% of LPS activated/engineered donor B cells acquired a germinal center (GC)-independent (CD73^-^) memory phenotype^27,28^ (Fig. 2**a-c**). This suggested engineered cells should be poised for successful vaccination at this timepoint.

**Figure 2.**
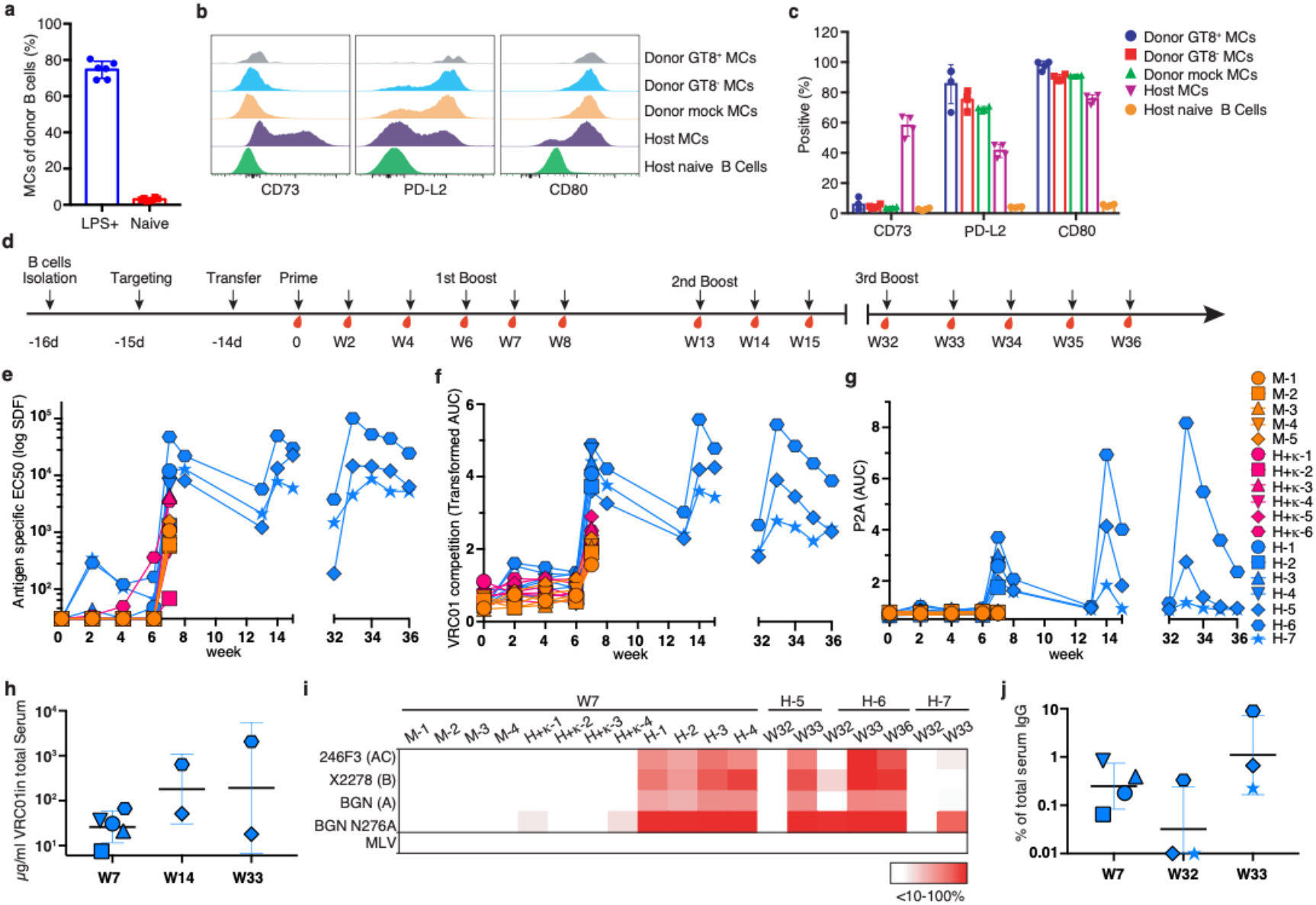
Engineered cell status after adoptive transfer and serological analysis of VRC01 responses after vaccination. **a, LPS activated donor cells acquire a memory phenotype *in vivo* after adoptive transfer.** Primary cells were either directly engrafted or cultured for 48 hrs in LPS before engraftment as required for B cell genome editing. The fraction of donor cells that showed a memory cell-like phenotype after 14 days in vivo are shown for n=6 animals in each group. Donor memory B cells are gated as Live, CD19^+^CD38^+^GL7^−^IgD^−^. **b, Engineered cells go to rest as germinal center independent memory cells (CD73^−^) *in vivo*.** Cells were targeted during *ex vivo* LPS culture and adoptively transferred into host animals. 14 days later successfully engineered (GT8^+^) cells were analyzed by flow cytometry. Host naïve and memory B cells are used as negative and positive controls. **c**, **Quantitation of B cells gated as in** (**b**). The fraction of successfully targeted cells with the indicated cell surface memory cell markers are given for n=4 engrafted animals. These are compared with unsuccessfully engineered cells in the targeted population (GT8^−^), mock engineered, or host cell controls. **d, Engineered B cell vaccine experimental design.** Time course of B cell engineering, cell transfer, immunization and blood draws in wild type C57BL/6J mice. (**e-g**) **Serum antibody responses after MD39-nanoparticle immunization in mice which received mock targeted, H-targeted, or H+κ targeted B cells.** Values indicate (**e**) serum dilution factor (SDF) EC50s of total (host+donor) response to immunogen. **f**, VRC01-competetive antibody responses indicated as area under the curve (AUC). **g**, Serum titers of Ig which carries the P2A tag from engineered L-chains, given as area under the curve (AUC). **h, μg/ml VRC01 in the serum.** The quantity of VRC01 in the indicated serum samples at the indicated timepoints was calculated by comparing serum dilution EC50 titers of the P2A peptide with that of a mouse VRC01-P2A tagged IgG standard. **i, serum neutralization of HIV.** The ability of IgG purified from the serum of the indicated animals at the indicated timepoints was tested for its ability to neutralize pseudovirus using the TZM-bl assay. Percent neutralization of virus achieved by 200μg/ml of IgG is given as a heat map for several tier-2 viruses from different clades (BGN=BG505). **j**, **VRC01 antibody as a fraction of the total serum IgG.** The VRC01 fraction of total serum IgG for the indicated animals at the indicated timepoints (weeks) was determined by comparing the BG505-N276A neutralization IC50 values for these samples with the human VRC01 IgG monoclonal antibody standard.

After targeting using either the *H* or *H+κ* engineering strategies, cells from cultures containing 40,000 VRC01 BCR-expressing cells were engrafted per animal into WT recipient mice. This corresponds to a VRC01-engineered B cell frequency in the host of approximately 3 in 100,000 B cells for animals containing either *H* or *H+κ targeted* cells (Fig. S2b). Mice were primed 14 days later (w0) with MD39-ferritin, a soluble stabilized HIV envelope native trimer immunogen multivalently presented on a ferritin-based nanoparticle which has a monomeric affinity for VRC01 of 124 nM^29,30^. Animals were boosted 6 weeks (w6) after a prime. Three mice engrafted with *H-targeted* cells involved in a longer-term study were further boosted 13 and 32 weeks after the prime (Fig. 2**d**). Total antigen-specific and engineered antibody responses were tracked in the serum by ELISA and using virus neutralization assays. Animals which received mock engineered or *H+κ targeted* cells elicited the lowest total antigen specific and VRC01-competitive antibody responses. Six of seven animals which received *H-targeted* cells showed on average 10-fold higher total antigen-specific titers 1 week after the first boost and this was accompanied by significantly higher levels of VRC01 competitive antibody in the serum (Fig. 2**e**,2**f**). Because some endogenous VRC01-competing antibody responses could be elicited by MD39 vaccination as illustrated in animals with mock-targeted cells, VRC01 in the serum of animals containing *H-targeted* cells was also measured by P2A ELISA, as VRC01 light chains expressed from these cells are tagged with a P2A self-cleaving peptide at the C terminus of the mouse kappa constant chain. Animals with *H-targeted* cells elicited P2A titers after each boost which could be quantified for high titer samples using a recombinant P2A-tagged VRC01 standard (Fig. 2**g**, 2**h**) The highest responder (H-6), produced ≈2 mg/ml of VRC01 in the serum by one week following the third boost. IgG purified from sera of these animals also showed HIV specific cross-clade neutralization of tier-2 viruses that bear hallmarks of typical circulating viruses such as relative neutralization resistance^31^ (Fig. 2**i**). Because no endogenous neutralizing responses could be elicited to a strain particularly sensitive to VRC01 neutralization (BG505-N276A), IC50s could be used to quantify VRC01 as a fraction of total serum IgG in responding animals (Fig 2**j**). In the best responder (H-6), VRC01 remained a significant fraction of the total IgG even after a 5-month rest period (≈0.33%), and this was boosted to ≈9% of the total IgG by one week after the third boost. We conclude that engineered B cells can make durable memory and antibody responses.

Consistent with the serum analysis, at 2 weeks after the first boost, animals that received *H-targeted* cells only showed high frequencies of VRC01-engineered GC B cells (Fig. 3**a-c**), splenic and bone marrow plasma cells (Fig. 3**d, e**) and MBCs (Fig. 3**f**). Engineered cells in animals from this group were expanded between 20-70-fold during the prime and boost (Fig. 3**g**) and VRC01 expressing memory (GT8^+^, KO11^−^, donor^+^) cells became mostly CD73^+^, implying their entry into GCs after vaccination unlike GT8^−^ donor cells which remained CD73^−^ after LPS treatment (Fig. 3**h**, 3**i**).

**Figure 3.**
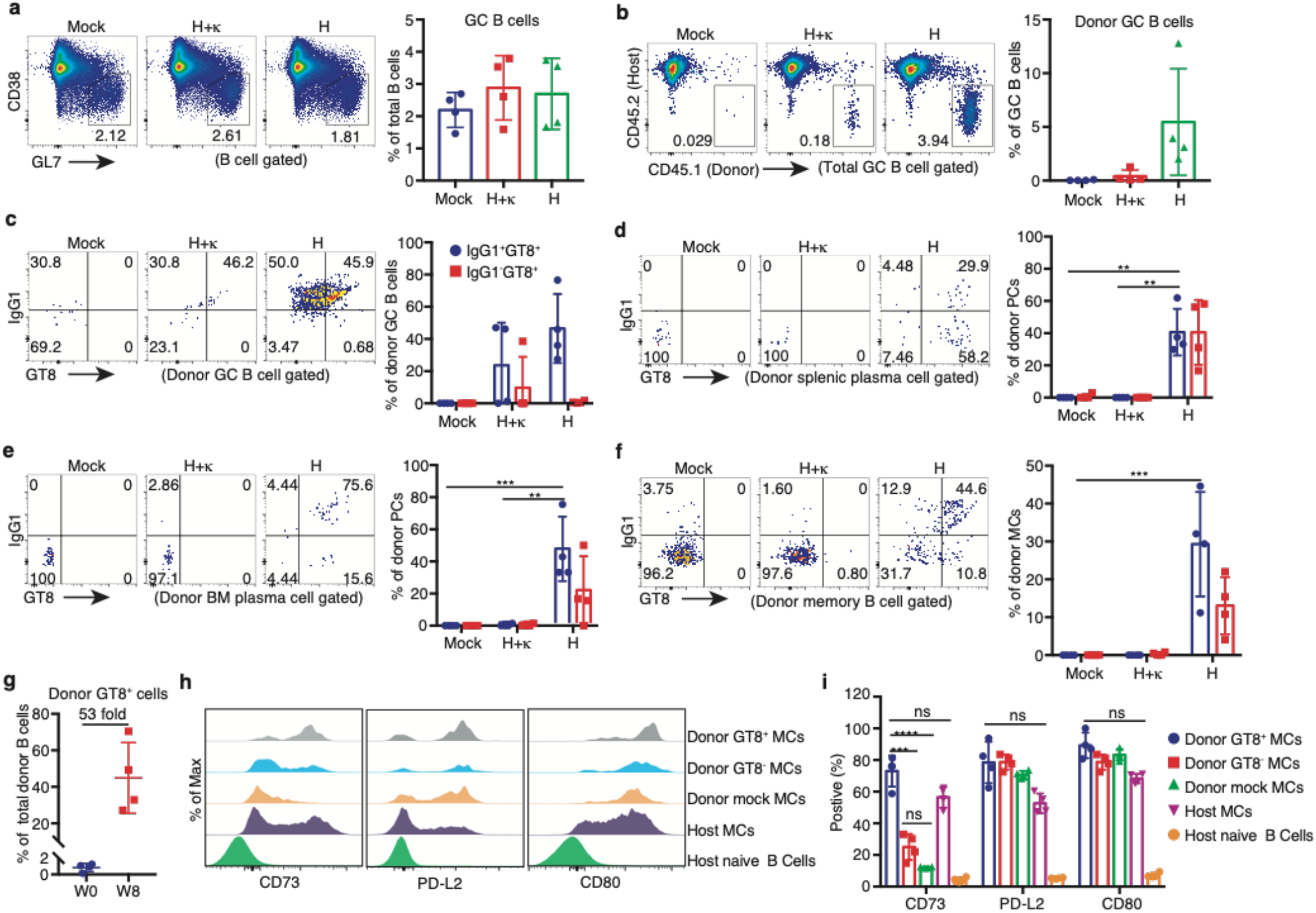
Flow cytometric analysis of antigen-matured engineered B cells. Host mice receiving LPS activated only (Mock), *H+κ*, or *H-targeted* B cells were analyzed 14d after the first boost (w8). n=4 animals in each group. **a-c, Germinal center B cells. a**, Representative flow cytometry and statistical analysis of total GC B cells pre-gated as live, CD19^+^ singlets. **b**, CD45.1^+^ (donor) GC B cells. **c**, Fraction of cells gated in **b** which were IgG^+^ (class switched) and GT8^+^KO11^−^ (engineered VRC01-expressing) **d-e, Plasma cells.** Frequency of class-switched and antigen-specific plasma cells (PCs) in spleen (**d**) and bone marrow (**e**) as a fraction of total donor PCs. GT8^+^KO11^−^CD45.1^+^CD45.2^−^IgG^+^ PCs were analyzed by intracellular staining after permeabilization of surfaced-stained cells. Plasma cells are gated as live, F4/80^−^IgD^−^TCRβ^−^ CD138^+^Sca-1^+^. **f, Memory cells.** Enumeration of IgG1^+^GT8^+^KO11^−^, engineered MCs (sIgD^−^ CD45.1^+^CD45.2^−^CD38^+^GL7^−^). **g**, **Vaccine induced expansion of engineered cells.** GT8^+^KO11^−^ CD45.1^+^CD45.2^−^ cell expansion at w8 compared with w0 as a percentage of the total donor cells. **h, Engineered memory cells have become germinal center dependent (CD73^+^) after vaccination.** GT8^+^ or GT8^−^ donor memory cells were compared with host naïve and memory B cells controls for expression of the indicated memory cell surface markers. **i, Quantification of B cells memory subtypes** gated as in **h. a-i**) Bars represent mean ± SD for all data points in each group. *, P< 0.05; **, P < 0.01; ns, not significant; unpaired 2-tailed T test.

Repeat experiments and variations on the original protocol also successfully elicited VRC01 memory responses in other groups of mice (Table S1). Variations included: 1) FACS enrichment of only antigen-specific engineered cells before transfer, 2) *ex vivo* activation of B cells using CD40L, IL-4, and CpG rather than LPS, 3) alternative immunization strategies in which engineered cells were primed coincident with transfer into mice with pre-existing immunogen-specific T cell help, and 4) immunization using different adjuvants.

Engineered immunoglobulin gene repertoires from the spleen (memory) and bone marrow (long-lived plasma cell) compartments were sequenced in several animals 42 days after their final boosts. Repertoires mostly conserved the original VRC01 amino acid sequence, however an abundance of variants with coding mutations were present with some highly mutated lineages emerging in the memory compartment (Figure 4**a**). While engineered cells were mostly IgM at time of transfer, the VRC01 repertoires after vaccination were almost completely class switched with IgG1 and IgG2 isoforms dominating as expected (Figure 4**a**). Coding mutations were observed across the entire length of the engineered VRC01 light chain and heavy chain variable regions, especially within the kappa constant gene, which is not normally in the path of the mutator (Fig. 4**b**). Coding changes in the P2A peptide were dramatically absent as this sequence must be highly conserved in order to express a functional BCR in engineered cells. Identical mutations could be observed across compartments and animals, and some coding changes appeared enriched in one compartment over the other within the same animal (Fig. 4**b**). The heavy chain CDR2 region of VRC01, which forms the primary interaction surface of this antibody with the virus^15^, showed coding changes after vaccination (Figure 4**c**) indicating that engineered responses generate some variation in antigen binding specificity which could be advantageous against highly diverse viral reservoirs and confirming the ability of engineered B cells to undergo somatic mutation and affinity maturation.

**Figure 4.**
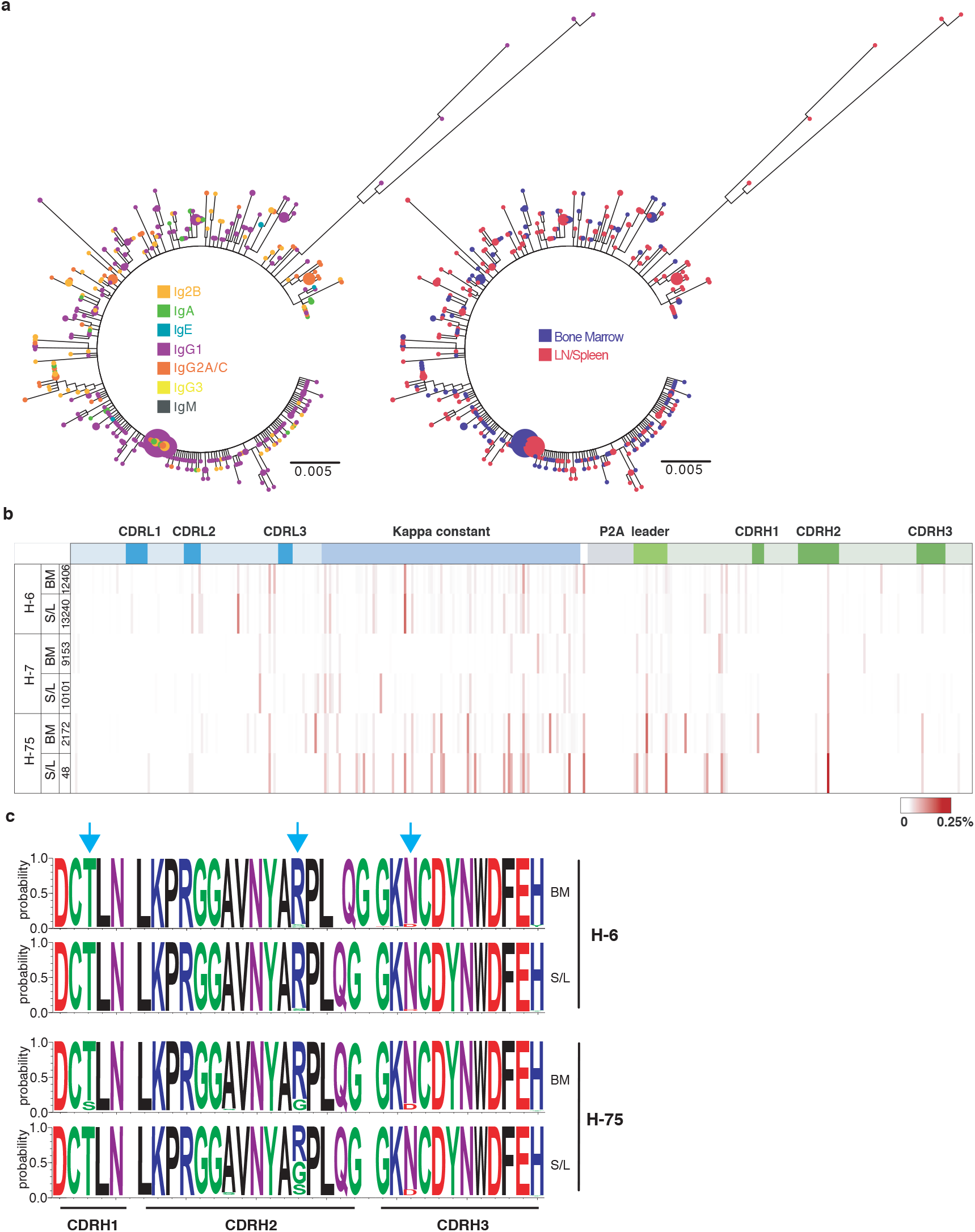
Engineered antibody repertoire in vaccinated animals. **a, Diversity of the engineered repertoire after immunization.** Relatedness of VRC01 clonotypes by isotype (left), and compartment (right), in one representative animal (H-7) 5 weeks after the final boost. **b, Mutational hotspots in the engineered VRC01 gene.** The frequency of amino acid changes at each residue position across the *H-targeted* VRC01 gene is shown as a percentage of the total sequences obtained for each dataset. 6 datasets are shown which are derived from both memory and plasma cell compartments from the three indicated animals. Specific coding changes across the length of the gene are given in Figure S3 from one representative animal (H-6). **c, Antigen binding properties are diversified in the engineered repertoire.** The fraction of sequences with coding changes from memory (S/L) or plasma cell (B/M) compartments are shown as sequence logos for the CDR heavy chain regions in the indicated animals. Blue arrows indicate amino acid positions undergoing diversification.

In the first demonstration of antibody activity, Behring and Kitasato showed in 1890 that serum from immunized animals could protect mice from lethal challenge of Diptheria and Tetanus toxins. Here we show that it is possible to passively transfer genetic information to the adaptive immune system that facilitates a high affinity, highly evolved, yet further evolvable, HIV bnAb response--in effect, demonstrating that one can program immune memory. Thus, we envision that our study represents a new phase in the development of passive immunity, started with the discovery of antibody itself.

## Materials and Methods

### Study design

Spleen-derived wild type mouse B cells can be activated and engineered *ex vivo* to express HIV broadly neutralizing antibodies as functional antigen receptors. The purpose of this study was to assess the ability of such engineered cells to expand and mature in response to HIV envelope-based vaccines in order to generate long-lived memory and high titer class-switched HIV bnAb responses in this wild type animal model.

### Cell culture

293T cells for HIV pseudovirus production were obtained from the American Type Culture Collection (ATCC, #CRL-3216) and cultured in DMEM (Invitrogen, #10313021) supplemented with; 10% fetal bovine serum (Invitrogen, #26140-079), 2mM L-glutamine (Invitrogen, #2503008), 100 units of Penicillin and 0.1 mg/ml of Streptomycin (Invitrogen, #15140122). TZM-bl cells for virus neutralization assays were obtained from the National Institutes of Health (NIH) AIDS Reagent Program (#8129) and cultured in DMEM supplemented with FBS (10%), 100 units of Penicillin and 0.1 mg/ml of Streptomycin. FreeStyle™ 293F cells for protein production were obtained from Life Technologies (#R79007) and cultured in FreeStyle™ 293 Expression Medium (Life Technologies, #12338018). Chemically competent DH5a *E. coli* for plasmid propagation were purchased from NEB (#C2987H).

### Animals

Animal studies were approved and carried out in accordance with protocols provided to the Institutional Animal Care and Use Committee (IACUC) at Scripps Research (La Jolla, CA) under approval number 18-0004. The mice were housed, immunized and bled at Scripps in compliance with the Animal Welfare Act and other federal statutes and regulations relating to animals in adherence to the Guide for the Care and Use of Laboratory Animals (National Research Council, 1996). All experiments were carried out in wild type 3-month old C57BL/6J (CD45.1/Ly5.1 or CD45.2/Ly5.2) male or female mice bred at Scripps Research Division of Animal Resources Facility (DAR). All procedures were performed on animals anesthetized using isoflurane.

### Cas9-RNP selection

gRNAs targeting the *IgH-J4* and *IgK-J5* region of the reference C57BL/6J genome (GRCm38) were designed using the GPP webtool (https://portals.broadinstitute.org/gpp/public/analysis-tools/sgrna-design). The gRNAs were synthesized by IDT (as crRNAs). CRISPR-Cas9 tracrRNA and HiFi Cas9 nuclease V3 were purchased from IDT (#1072534, #1081061). Cas9:crRNA:tracrRNA complexes were made according to manufacturer’s instructions. The targeting efficiency of different gRNAs was directly assessed by VRC01 engineering efficiency in LPS-activated primary B-cells (described below) and the best gRNAs were chosen for *IgH-J4* and Igκ- J5 targeting.

### HIV bnAb donor DNA plasmid preparation

#### H-targeting donor plasmids

VRC01 donor DNA was designed as follows from 5’ to 3’: 1) 498-basepair (bp) homology arm upstream of the mouse *IgH-J4* (in germline configuration). 2) The mouse *IgHV1-82* heavy chain promoter region (entire 5’UTR). 3) The mouse *IgKV4-53* leader (intron removed) and human VRC01 light-chain variable gene followed by the mouse kappa constant region without stop codon. 4) A GSG linker followed by the P2A self-cleaving peptide sequence. 5) The mouse *IgHV1-82* leader lacking the intron followed by the human VRC01 heavy-chain variable gene and constant gene donor splice site. 6) A 500-bp region of DNA homologous to the *IgH-J4* intron beginning after *JH4*. The homology regions and V-gene promoter were amplified from C57BL/6J gDNA. The VRC01 LC-P2A-HC VDJ gene was synthesized (Geneart).

#### H+κ targeting donor plasmids

The *H-targeting* plasmid was modified by deletion of the LC and P2A peptide and adding in the IgHV1-82 intron. The LC donor DNA was designed as follow from 5’ to3’: 1) 510bp homology arm upstream of the mouse *IgK-J5;* 2) the mouse *IgKV4-53* light chain promoter (entire 5’ UTR), leader and V-gene intron; 3) VRC01 VJ and constant gene donor splice site 4) a 510bp region downstream of *Jκ5*. All fragments were assembled using Gibson assembly (NEB, #E5510S) according to the manufacturer’s instructions into the PICOZ (PMID: 31124712) carrier vector synthesized from IDT and transformed into DH5a *E. coli*. (NEB, C2987H). Single colonies from bacteria plated on agar containing Zeocin were cultured; plasmids were isolated (Qiagen, #27106) and sequenced (Eton Biosciences) using several primers to generate high quality coverage of the entire donor DNA region. Donor DNA plasmids were purified using EndoFree plasmid maxi kit (Qiagen, #12362) for engineering experiments.

### B-cell activation and engineering

All work was conducted under sterile conditions. B-cells isolated from the spleens of CD45.1 or CD45.2 congenic mice using negative selection (Miltenyi, #130-090-862) were cultured in RPMI-1640 (Invitrogen, #21870076) supplemented with 1X NEAA (Invitrogen, #11140050), 1X sodium pyruvate (Invitrogen, #11360070), 55μM 2-Me (Invitrogen, #21985023), 10% FBS (Invitrogen, #26140-079) and either 1) 50 μg/ml LPS (Sigma, #L2880-100MG) or 2) CpG ODN 2006 at 1μM (Synthesized from IDT), IL-4 at 50ng/ml(Biolegend, #574306), anti-CD40 at 5 μg/ml (Thermo, #16-0401-86) for 30 hours. The Alt-R CRISPR-Cas9 system components (IDT) were used to make Cas9-RNP complexes according to the manufacturer’s instructions. gRNA sequences for *H* or *κ* targeting were 5’-gagaggccattcttacctg-3’ and 5’-ttacgtttcagctccagct-3’ respectively. Activated cells were washed three times with 1X DPBS (Invitrogen, #14190-144) and resuspended at 5 million cells/ 100 μl in Neon R buffer (Invitrogen, #MPK10096). For H-targeting, 5 μl H-targeting Cas9-RNP complexes, 2.16 μl 100 μM CRISPR electroporation enhancer (IDT), and 20 μg donor DNA (4 μl) were mixed and added to the resuspended cells. For *H*+*κ* -targeting, 5μl each of *H*- and *κ*-targeting Cas9-RNP complexes, 4.32 μl 100 μM CRISPR electroporation enhancer (IDT), and 15 μg each of heavy chain and light chain donor DNAs were mixed and added to the resuspended cells. The cells were electroporated using a 100 μl tip (Invitrogen, MPK10096) with 1650 V, 20 ms, 1 pulse. Nucleofected cells were immediately cultured in media without cell activation components for 1 h before these components were added for further *ex vivo* culture.

### Adoptive transfer of engineered cells

18 h after nucleofection, activated mock or engineered cells were washed 4 times with 1X DPBS (Invitrogen, #14190-144) for adoptive transfer into recipient animals. Some cells were further cultured to confirm the quantity of successfully targeted cells by flow cytometry 48 hrs post nucleofection. VRC01-engineered B cells were identified by staining for viable (propidium iodide negative), eOD-GT8-positive, eOD-GT8-KO11-negative cells. In some experiments (Table S1), successfully targeted cells were enriched by FACS before transfer 18h post-nucleofection by gating the live, GT8^+^KO11^−^ population.

### Immunization of rested cells

4 million *H+κ targeted* VRC01 cells or 1 million *H-targeted* VRC01 cells (corresponding to 40,000 VRC01-expressing engineered cells by either strategy) were retro-orbitally transferred into recipient animals. These animals were rested for 14 d before immunization with a 200 μl mixture of 20 μgs of immunogen and 1) Ribi adjuvant (according to the manufacturer’s instructions, Sigma, #S6322-1VL) or 2) 5 μgs of IscoMPLA (kindly provided by the lab of Darrell Irvine at MIT) in 1x DPBS administered by i.p. injection. Mice were boosted with the same immunogen 6, 13 and 32 weeks after priming. Serum samples were collected through ocular bleeding of animals at time intervals indicated in Figure 2. All procedures were done on isoflurane anesthetized mice.

### Immunization of T-primed cells (Table S1)

Recipient animals were i.p. immunized with 200 μl mixture of 10 μg of either the lumazine synthase or ferritin base in 1X DPBS and Ribi adjuvant. 7 days later, targeted cells containing 65,000 VRC01-expressing engineered cells were retro-orbitally transferred into pre-primed host animals. Lumazine synthase pre-primed animals were immunized with 20 μg eOD-GT8-60mer and ferritin pre-primed animals were given 20 μg MD39-8mer immunogen in Ribi adjuvant at the same time. For the FACs enriched group, 300,000 sorted antigen specific cells were transferred into recipient mice following the same manner of T-primed immunization. All procedures were done on isoflurane anesthetized mice.

### Flow cytometry analysis and single-cell sorting

Spleen suspensions were generated by smashing the spleen between frosted glass slides. Bone marrow was released from tissue-free tibia and femurs by a mortar-and pestle. Red blood cells were lysed with ammonium chloride (0.83%) before filtering cells through a 40 μM cell strainer to generate single-cell suspensions. Fc Blocker (homemade mAb 2G4) was added to single-cell suspensions at 0.5 μg per 10^6^ cells before antibody staining. For antigen-specific analysis, biotinylated AviTagged GT8 monomers were pre-complexed for 30 min to Streptavidin-AF488 (Thermo, #S32354) or Streptavidin-AF647 (Thermo, #S32357); GT8-KO11 was complexed to streptavidin-BV421 (BD, #BDB563259). Fluorophore-conjugated antibodies CD45.1 (Biolegend, #110728), CD45.2 (Biolegend, #109806), GL7 (Biolegend, #144608, #144610) TCRb (Biolegend, #109228), F4/80 (Biolegend, #123128), Ter119 (Biolegend, #116228), CD38 (Biolegend, #102718), IgD (Biolegend, #405710), IgM (Biolegend, #406512), IgG1 (Biolegend, #406620), CD138 (Biolegend, #142504), Sca-1 (Ly6A/E) (Biolegend, #122512) and CD19 (Biolegend, #152408), CD80 (Biolegend, #104712), CD73 (Biolegend, #127210), PD-L2 (Biolegend, #107216), anti-mouse Kappa (Clone 187.1 AF647), Anti mouse Lambda (Biolegend, #407306) were used to define different cellular populations. CD45.1 and CD45.2 mAbs were used to distinguish host and transferred B-cells. IgG1 staining was used to identify class switched B-cells. VRC01 expressing memory cells (MCs) were gated as live GT8^+^KO11^−^CD19^+^CD38^high^sIgD^−^GL7^−^; Germinal center (GC) B-cells were gated as live CD19^+^ CD38^−^ GL7^+^; Plasma cells (PCs) were gated as CD138^+^ Sca1^+^TCRb^−^Ter119^−^sIgD^−^sIgM^−^GL7^−^F4/80^−^; Permeabilized surface-stained PCs were intracellularly stained with GT8, KO11 and IgG1 probes. Cells were analyzed using the Cytek Aurora or sorted using a BD FACSAria at the Scripps Flow core.

### Peripheral tolerance

8 million B cells isolated from the spleens of CD45.2 congenic mice using negative selection (Miltenyi, #130-090-862) were either transferred directly or after 24 hours of ex vivo LPS-activation into CD45.1 pAlb and WT recipient mice at day 0. Before adoptive transfer, the frequency of B cells carrying kappa light chain were determined using an AF647-labelled anti-kappa antibody (homemade) by flow cytometry. The Kappa frequency of CD45.2 (donor) from splenic B cells of recipient mice were analyzed successively on day 1, 8, 15 and 28 post transfer.

### Engineered B cell Immunoglobulin Repertoire Sequencing and Analysis

Cells were released from mouse spleen and lymph nodes by pressing these tissues through a 0.2 μm cell strainer using a rubber syringe plunger and rinsing them into a 50 ml falcon tube using MACS buffer (PBS+2%FBS+2mM EDTA). Bone marrow was obtained by crushing tissue-free tibia and femur bones with a mortar and pedestal and rinsing the released cells through a 0.2 μm cell strainer into a 50 ml falcon tube. Cells were pelleted by centrifugation (600 x g for 6 min). Red blood cells (RBCs) were disrupted by resuspending the cell pellets in 10 ml of RBC lysis buffer (155mM NH_4_Cl + 12mM NaHCO_3_ + 0.1mM EDTA) for 3 min at RT. Cells were then diluted to 50 ml with MACS buffer and pelleted. The cells were washed with 50 ml of MACS buffer and cell numbers determined using a Coulter Particle Counter. We routinely obtained approximately 1-1.4 × 10^8^ cells from the spleen and lymph node samples, and roughly 6-9 ×10^7^ cells from the marrow. Cells were pelleted and resuspended in 100 μl MACS buffer/10^7^cells. 0.5 μg of FcR blocking reagent was added per 10^6^ cells and the mixture incubated for 10 min at RT. 10^6^ cells were kept for FACS analysis. The remaining cells were then labelled with biotinylated anti-CD45.2 Ab (Clone 104, Biolegend, #109804) at a 1:100 dilution for 30 min. The cells were then washed twice with 1-2 ml MACS buffer per 10^7^ cells. Cell pellets were resuspended in 70 μl MACS buffer and 20 μl of anti-biotin microbeads (Miltenyi, #130-090-485)/10^7^ cells. The suspension was well mixed by pipetting and incubated for 15 min at 4°C. The cells were washed by adding 1-2 ml of MACS buffer per 10^7^ cells and pelleted. Cells were then resuspended in 500μl of MACS buffer and pipetted onto a MACS buffer rinsed LS column (Miltenyi, #130-042-401) placed in a magnet field to retain labelled cells on the column. The column was rinsed 3× 3 ml MACS buffer. The column was then removed from the magnet and labelled cells eluted into a flacon tube with 7 ml MACS buffer pushed through the column with a plunger. These cells were then subjected to a second round of purification over a new LS column. Cell numbers were measured and cells pelleted for mRNA purification. We routinely obtained approximately 4-6 × 10^6^ and 2-4 × 10^6^ cells for spleen/lymph node or bone marrow derived cells respectively. 10^5^ cells were kept for FACS analysis. mRNA was purified using the Qiagen RNeasy micro kit (Qiagen, #74004) according to the manufacturer’s instructions. mRNA was immediately subjected to reverse transcription using the Superscript III first strand synthesis kit (Thermo) to generate cDNA following the manufacturer’s instructions. For the RT-PCR we used either a VRC01-specific primer (5’-CCATCTCATCCCTGCGTGTCTCCGAC NNNNNNNN GATGAGACGATGACCG-3’) or a mixture of isotype (IgM, IgA, IgE, IgD & IgG) specific primers (PID_mIgM-CCATCTCATCCCTGCGTGTCTCCGAC NNNNNNNN CTGGATGA CTTCAGTGTTGT; PID_mIgA-CCATCTCATCCCTGCGTGTCTCCGAC NNNNNNNN CCAGGT CACATTCATCGTG; PID_mIgE-CCATCTCATCCCTGCGTGTCTCCGAC NNNNNNNN GTTCA CGTGCTCATGTTC; PID_mIgD-CCATCTCATCCCTGCGTGTCTCCGAC NNNNNNNN GCCAT TTCTCATTTCAGAGG; PID_mIgG12- CCATCTCATCCCTGCGTGTCTCCGAC NNNNNNNN KK ACAGTCACTGAGCTGCT; PID_mIgG3- CCATCTCATCCCTGCGTGTCTCCGAC NNNNNNNN GTACAGTCACCAAGCTGCT) containing unique primer IDs (PID) and a primer landing site. All primers for cDNA synthesis were HPLC purified. The primer IDs were tagging each RNA template with a unique 8-nucleotide-long identifier, which allowed us to group amplified sequences for each individual template. The primer landing site was used for the reverse primer binding site of the subsequent hemi-nested PCRs. The resulting cDNA was purified using AMpureXP beads (Beckman Coulter, #A63882) at a volume ratio of 1 : 1. The purified cDNA was amplified and barcoded during a hemi-nested PCR using the forward primers VRC01_L1-F (5’– GGATTTTCATGTGCAGATTTTCAGCTTCATGC -3’) for the 1^st^ round and the primer VRC01_Junc-F (5’- CAGTGTCACAGTCATATTGTCCAGTGG -3’) for the 2^nd^ round PCR in combination with the reverse primer PID-R (5’-ATCCCTGCGTGTCTCCGAC -3’), that attached to the primer landing site. Successful amplification was confirmed on a 0.7% agarose gel and amplicons were barcoded during an additional 2^nd^ round PCR using a barcoded 2^nd^ round primer set. The final barcoded amplicons were quantified using the Tapestation D5000 ScreenTape System (Agilent) and pooled at equimolar ratios. Library preparation and sequencing of SMRTbell template libraries of approximately 1.5-kb insert size were performed according to the manufacturer’s instructions (Pacific Biosciences).

Data was processed and visualized using a pipeline built for this sequencing protocol in the Julia language, using NextGenSeqUtils ^1^. Briefly, samples were first demultiplexed and oriented using demux_dict(), and reads were dereplicated. Sequences were collapsed by primer ID (UMI), and if sequences with the same PID differed, either due to sequencing error or PID “clashes”^2^ we retained the most frequent variant. This gives us a dataset at the “transcript level”. To exclude PacBio or RT indel errors, we discard reads with any indel variation relative to the engineered reference. For isotype mixture RT sequencing, we perform isotype calls on transcripts by matching the sequence immediately 3’ of the engineered polypeptide against isotype references extracted from the CH1 IMGT database, discarding transcripts that do not match, or match ambiguously. We then collapse “transcripts” into “variants” (sequences with 100% identical nucleotides over the polypeptide region, and, in the isotype RT datasets, identical isotype calls), retaining a transcript count for each variant. These variants, and their associated frequencies, were used in all sequence analyses (always including the variant frequency when counting any mutations). For phylogenetic analysis, but nowhere else, we additionally collapse any “singleton” variants (where frequency = 1) into larger variants of the same isotype that are just one nucleotide distance from them (we retain singletons that are >1 nucleotide from their nearest variant). This allows for compact display of phylogenies. Maximum likelihood phylogenetic trees were inferred using FastTree2^3^, and visualized using FigTree (http://tree.bio.ed.ac.uk/software/figtree/).

### Total response ELISA

384-well ELISA plates (Corning, #3700) were initially coated with 12.4 μl/well streptavidin (Jackson Immuno Research Labs, #016-000-084) at 2 μg/ml diluted in PBS and incubated at 4°C overnight. Plates were washed 3x with 100 μl/well PBS containing 0.05% Tween (PBS-T) and blocked with 40 μl/well PBS + 3% BSA at RT for 1 hour before the addition of 12.4 μl/well biotin-labeled BG505 SOSIP (produced in house, Pugach et al., *Journal of Virology*, 2015) at 2 μg/ml diluted in PBS-T and 1% BSA. Mouse sera were serially diluted (3x) using PBS-T and 1% BSA starting at 1:10 and 12.4 μl/well incubated at room temperature for 1 hour. After washing (as above), 12.4 μl/well alkaline phosphatase-conjugated goat anti-mouse IgG (H+L) (Jackson Immuno Research Labs, #115-055-146) diluted 1:5000 in PBS-T, 1% BSA was added and incubated for 30 min at room temperature (RT). Plates were washed and p-Nitrophenyl Phosphate (pNPP) substrate (Sigma Aldrich, #S0942) dissolved to 1 mg/mL in substrate buffer (10 mM MgCl_2_ with 80 mM Na_2_CO_3_ and 15 mM NaN_3_, pH 9.8), was added at 12.4 μl/well to visualize the binding of antigen specific mouse IgG. Optical density (OD) at 405 nm was read on a Molecular Devices (SpectraMax Plus) plate reader, allowing the same amount of development time for each plate. EC_50_ values were generated by fitting curves to plots of Absorbance values vs. the log of the serum dilution for each sample.

### Anti-P2A ELISA

384-well plates were pre-coated overnight at 4°C with 12.4 μl/well, 2 μg/ml BG505 SOSIP 2JD6 nanoparticle produced in house^4^. Plates were washed and blocked with 40 μl/well of PBS supplemented with 3% BSA at RT for 1 hr and washed again. Mouse serum samples serially diluted (2x) with PBS-T and 1% BSA starting at 1:10 dilution were added (12.4 μl/ well) and incubated at RT for 1 hour. A P2A-light chain tagged mouse VRC01 IgG monoclonal antibody standard starting at 1 μg/mL was also added (12.4 μl), serially diluted (2x), and incubated at RT for 1 hour. Plates were washed and incubated with 12.4 μl/well biotin-labeled anti-2A peptide (3H4) mouse antibody (NovusBio, #NBP2-59627) at 1 μg/ml in PBS-T and 1% BSA. Plates were washed and the captured complex incubated with 12.4 μl/well alkaline phosphatase conjugated Streptavidin (Jackson Immuno Research Labs, #016-050-084) at 1:3000 dilution in PBS-T and 1%BSA at RT for 1 hr. Plates were washed and pNPP substrate was added as above. All plates developed for the same period of time before being read at 405 nm as above. P2A titers were reported as area under the curve for plots of Absorbance vs. log dilution factor for each sample. When maximum Abs 405nm values were above 1.5, P2A-antibody quantities in serum samples were calculated by multiplying the sample EC50 with that of the mouse-VRC01 standard.

### Mouse VRC01-P2A IgG Preparation

The VRC01 LC-P2A-HC construct was PCR-amplified from engineered C57BL/6J mouse cDNA and cloned between the promoter and constant regions of a mouse IgG2b heavy chain expression vector (InvivoGen, #pfuse-mchg2b) using Gibson Assembly. A leader sequence (5’ATGGGATGG TCATGTATCATCCTTTTTCTAGTAGCAACTGCAACCGGTGTACATTCA3’) was also incorporated 5’ to the LC region. Transient expression of the P2A-VRC01 antibody was accomplished through transfection into FreeStyle™ 293-F cells following manufacturer’s guidelines. The supernatant was harvested 5 days post transfection and IgG was affinity purified using Protein A Sepharose (GE Healthcare, #17-5280-02). SDS-PAGE was used to confirm the purity of IgG and adequate cleavage of the P2A peptide. Activity of the antibody was comparable to human VRC01 IgG monoclonal antibody in virus neutralization assays.

### Competition ELISA

384-well plates were pre-coated overnight at 4°C with 12.4 μl of BG505 SOSIP 2JD6 at 2 μg/ml diluted in PBS. After washing and blocking, 12.4 μl/well of mouse sera serially diluted (2x) in PBS-T and 1% BSA (starting at 1:10) was added to plates and incubated at RT for 1 hour. 12.4 μl/well of human VRC01 monoclonal antibody at 200 ng/ml was then added directly to the diluted sera. Plates were incubated for another hour after mixing and non-binding antibodies were washed away. Captured antibodies were detected with alkaline phosphatase-conjugated goat anti-human IgG, Fc_γ_ fragment specific (Jackson Immuno Research Labs, #109-055-098) diluted 1:5000 in PBS-T and 1% BSA. Plates were incubated at RT for 1 hour before the addition of pNPP substrate as described above. OD values at 405 nm were read, and standard curves were generated. Absorbance values were transformed by taking the absolute value of the (sample Abs405nm minus the Abs405nm of non-competing negative control wells). The area under the curve was then generated from Abs vs log serum dilution plots.

### Neutralization assay

Under sterile BSL2/3 conditions, PSG3 ^5^ plasmid was co-transfected into 293T cells along with various HIV envelope plasmids ^6^ using Lipofectamine 2000 transfection reagent (ThermoFisher Scientific, #11668019) to produce single-round of infection competent pseudo-viruses representing multiple clades of HIV. 293T cells were plated in advance overnight with DMEM medium +10% FBS + 1% Pen/Strep + 1% L-glutamine. Transfection was done with Opti-MEM transfection medium (Gibco, #31985) using Lipofectamine 2000. Fresh medium was added 12 hours after transfection. Supernatants containing the viruses were harvested 72h later. In sterile 96-well plates, 20 μl of virus was immediately mixed with 20 μl of serially diluted (3x) purified IgG from mouse sera (starting at 400 μg/ml) and incubated for one hour at 37°C to allow for antibody neutralization of the pseudoviruses. 5,000 TZM-bl cells/ well (in 40 μl of media containing 100 μg/ml Dextran) were directly added to the antibody virus mixture. Plates were incubated at 37°C for 48 h. Following the infection, TZM-bl cells were lysed using 1X luciferase lysis buffer (25mM Gly-Gly pH 7.8, 15mM MgSO_4_, 4mM EGTA, 1% Triton X-100). Neutralizing ability disproportionate with luciferase intensity was then read on a Luminometer with luciferase substrate according to the manufacturer’s instructions (Promega, #PR-E2620).

### Statistical analysis

Statistical analysis used an unpaired two-tailed T test (Prism, Graphpad). Correlation between EC50 of P2A titers and the relative concentration of VRC01 was calculated using Pearson correlation coefficient with linear regression (Prism, Graphpad). *P < 0.05, **P < 0.01, ***P < 0.001, ****P < 0.0001

## Acknowledgements

We thank Nicolle Jigarjian and staff at the Scripps Department of Animal Resources for ongoing care of experimental animals, Darrell Irvine at MIT for providing IscoMPLA adjuvant, and Christina Corbaci for assistance making figures. This work was supported by the Bill and Melinda Gates Foundation (grant number OPP1183956 to J.E.V) and by the National Institutes of Health (5R01DE025167-05 to D.R.B. and R01AI128836 and R01AI073148 to D.N.).

## Author Contributions

D.H. and J.T.T. developed methods and reagents for B cell engineering, carried out adoptive transfer and immunization experiments, collected animal serum and tissues, performed ELISA and FACS analysis. A.O. performed ELISA, virus neutralization assays and produced immunogens and antibodies. T.V, M.V.G and B.M. generated engineered repertoire libraries, sequenced the libraries, analyzed the sequence datasets, generated figures and edited the manuscript. M.T., J.A., L.C., T.R.B., K.S., W.L., E.L., A.G.M. assisted with methods development, experiments or animal care. T.S. and W.S. provided HIV vaccine immunogens and FACS probes. J.E.V., D.H. and D.N. designed the experiments. J.E.V., D.N., D.H. and D.R.B analyzed the data and wrote the manuscript.

---

**Figure S1.**
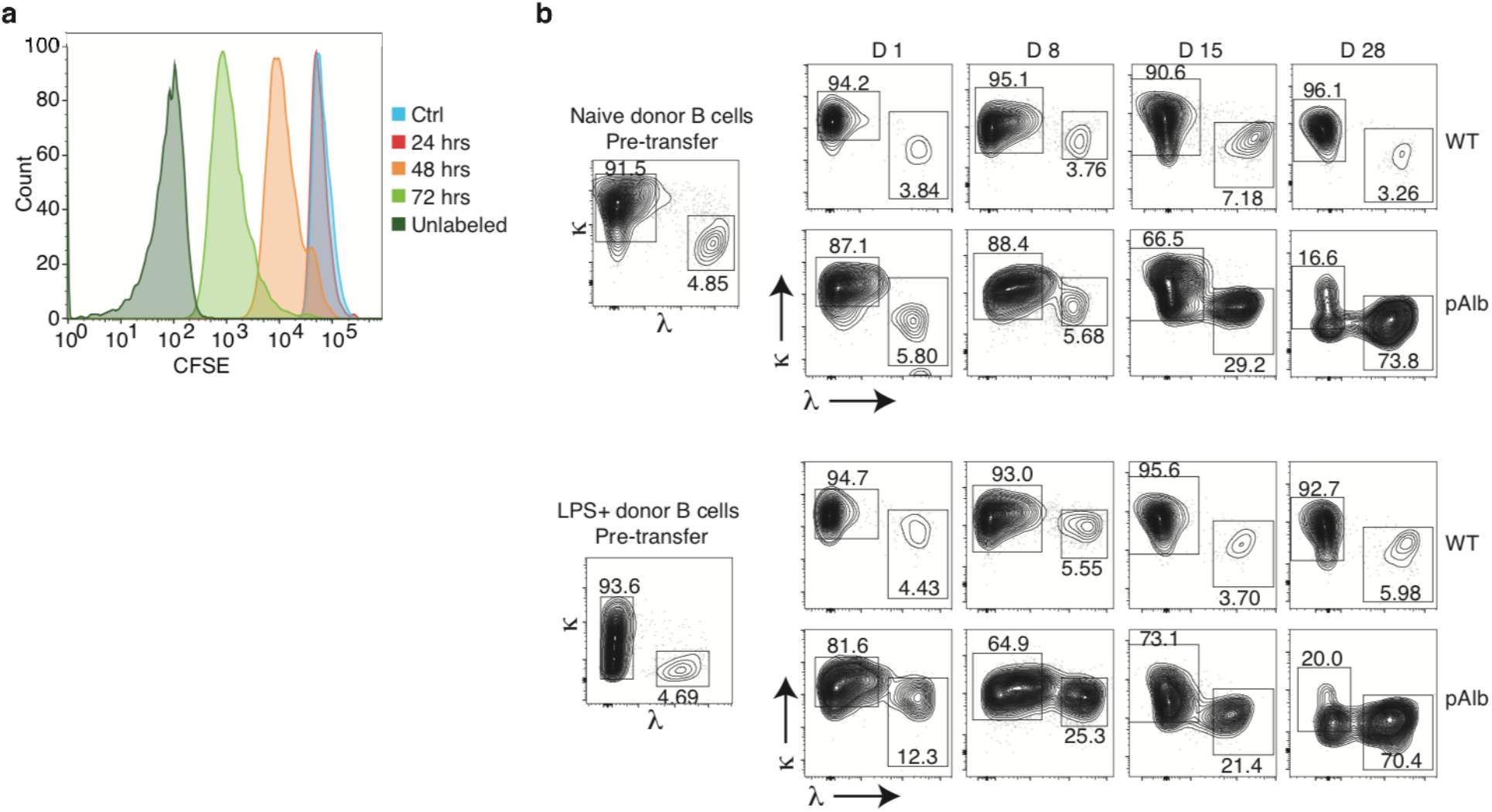
LPS culture and peripheral tolerance. **a**, B cells were isolated, labeled with CFSE, stimulated *in vitro* with LPS (50 μg/ml) and assessed for proliferation at the indicated times by flow cytometry. **b**, A representative gating strategy for donor kappa and lambda B cells. Naïve and LPS activated B cells were transferred to pAlb mice expressing anti-kappa superantigen in the liver. Splenocytes from host mice 15d post transfer were analyzed by flow cytometry to detect the kinetic changes of donor kappa and lambda B cells. Statistical analysis of kappa kinetic changes is shown in Figure 1d.

**Figure S2.**
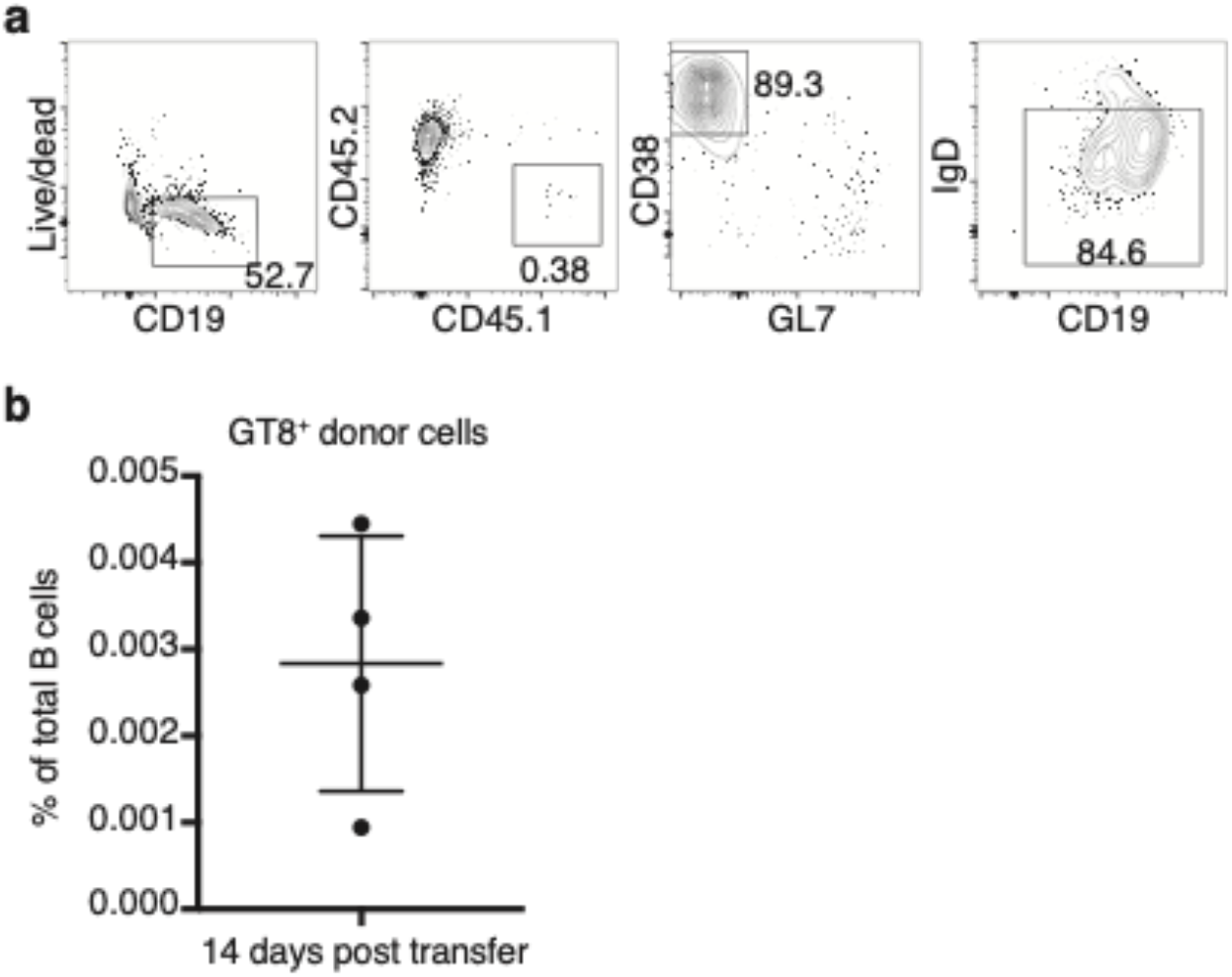
Flow cytometry analysis of adoptive transferred engineered B cells. **a**, A representative gating strategy for memory B cells. Naïve or LPS-activated engineered B cells were transferred to WT immunocompetent mice. Host splenocytes are analyzed by flow cytometry 14d post transfer to identify the status of donor B cells. Splenocytes from mice receiving LPS-activated engineered B cells were chosen to represent memory B cells gating strategy. Memory B cells were gated as CD19^+^CD38^+^GL7^−^IgD^−^. Donor and host memory B cells were further gated as CD45.1^+^ and CD45.2^+^, respectively. Engineered antigen-specific B cells were gated as CD45.1^+^GT8^+^KO11^−^ memory B cells. **b**, Statistical analysis of the frequency of engineered VRC01 cells among splenocytes 14d post transfer.

**Figure S3.**
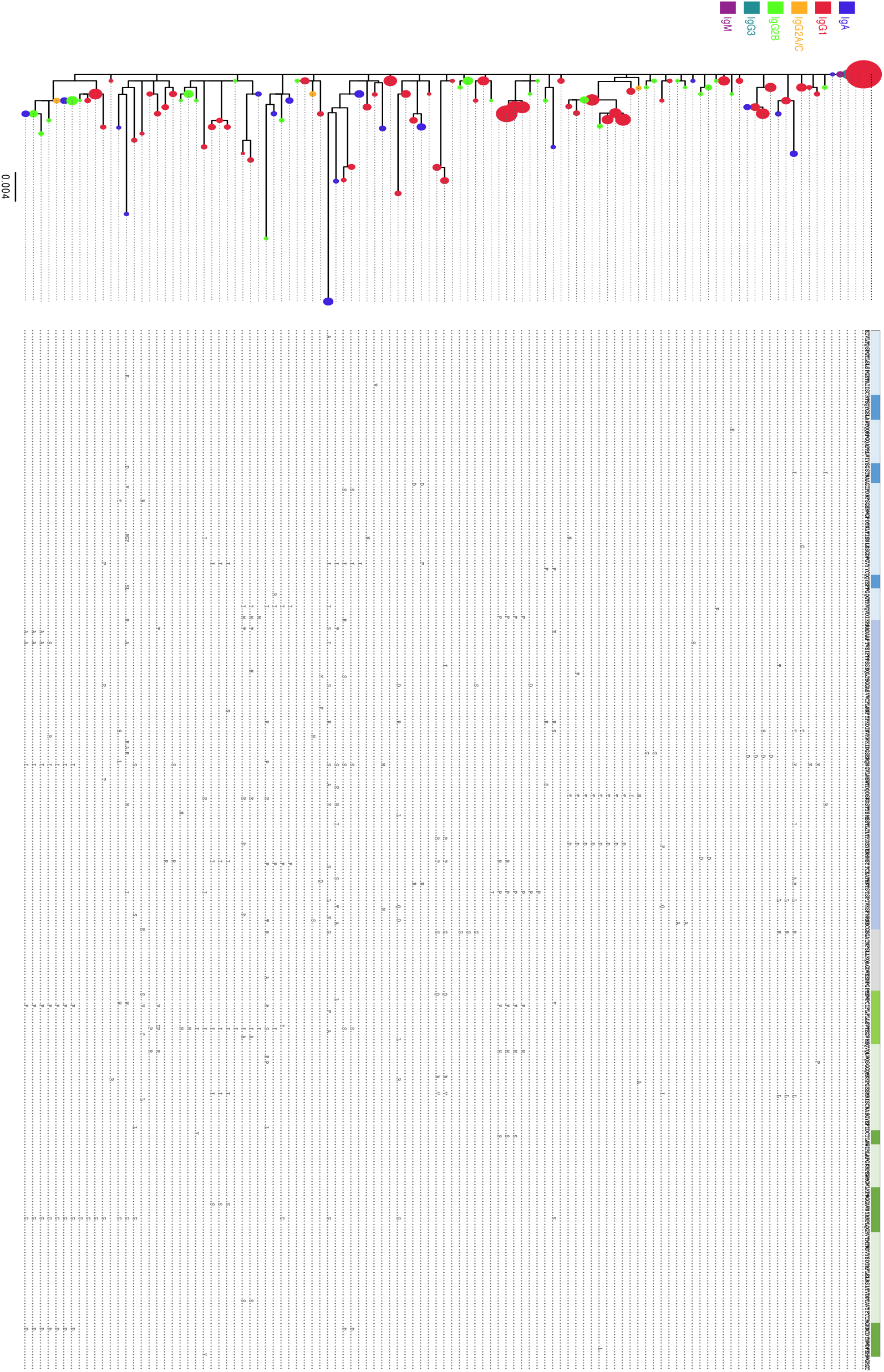
VRC01 repertoire sequence alignment from a boosted animal. Coding sequence changes are shown along the length of the VRC01 gene corresponding to the indicated clonotype.

**Table S1:**
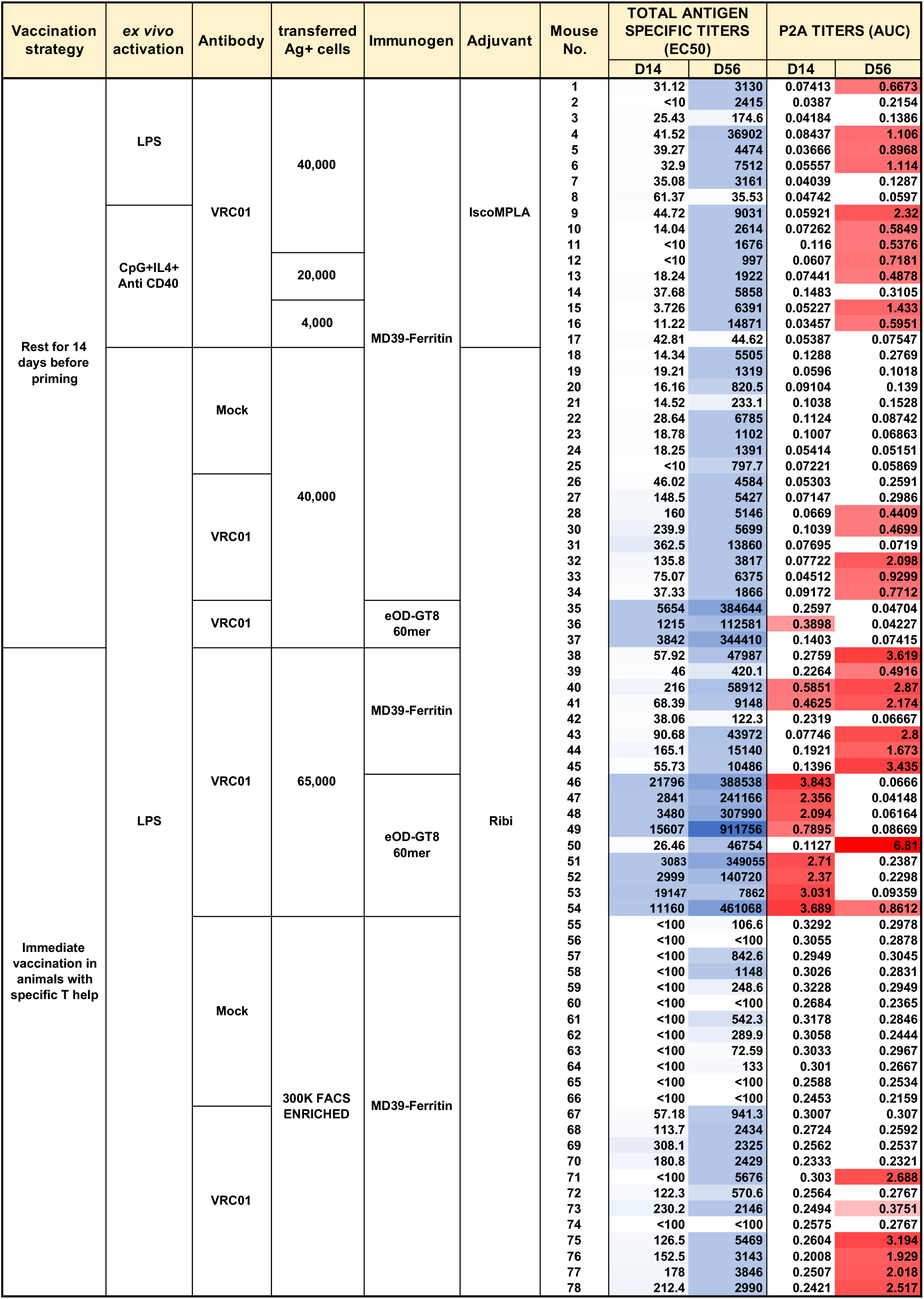
Engineered B cell vaccine results in other animals. Reproduction or variation of vaccination experiments are shown. Parameters varied is to the left of the animal number. Total antigen specific or engineered (P2A) Ab titers elicited 2 weeks after prime and 2 weeks after boost are shown on the right.

